# Comprehensive analysis of retinal development at single cell resolution identifies NFI factors as essential for mitotic exit and specification of late-born cells

**DOI:** 10.1101/378950

**Authors:** Brian S. Clark, Genevieve L. Stein-O’Brien, Fion Shiau, Gabrielle H. Cannon, Emily Davis, Thomas Sherman, Fatemeh Rajaii, Rebecca E. James-Esposito, Richard M. Gronostajski, Elana J. Fertig, Loyal A. Goff, Seth Blackshaw

## Abstract

Precise temporal control of gene expression in neuronal progenitors is necessary for correct regulation of neurogenesis and cell fate specification. However, the extensive cellular heterogeneity of the developing CNS has posed a major obstacle to identifying the gene regulatory networks that control these processes. To address this, we used single cell RNA-sequencing to profile ten developmental stages encompassing the full course of retinal neurogenesis. This allowed us to comprehensively characterize changes in gene expression that occur during initiation of neurogenesis, changes in developmental competence, and specification and differentiation of each of the major retinal cell types. These data identify transitions in gene expression between early and late-stage retinal progenitors, as well as a classification of neurogenic progenitors. We identify here the NFI family of transcription factors (*Nfia*, *Nfib*, and *Nfix*) as genes with enriched expression within late RPCs, and show they are regulators of bipolar interneuron and Müller glia specification and the control of proliferative quiescence.

## INTRODUCTION

Neural progenitor cells (NPCs) of the developing central nervous system (CNS) undergo a series of stereotypical, stage-dependent transitions during the course of neurogenesis. These include an initial transition from a slowly proliferating, relatively quiescent neuroepithelial status to actively proliferating neurogenic progenitors (Martynoga *et al.*, 2012; Schmidt *et al.*, 2013); a progressive transition of NPCs from symmetric proliferative to asymmetric self-renewing divisions, and later to terminal neurogenic divisions (Homem *et al.*, 2015; Taverna *et al.*, 2014); and changes in developmental competence - the ability of NPCs to give rise to different subtypes of neurons and glia (Cayouette *et al.*, 2013; Kohwi and Doe, 2013; Okano and Temple, 2009). While the molecular mechanisms that control these temporally dynamic processes in the vertebrate nervous system are still largely not understood, the retina offers a relatively tractable system for these studies, owing to its relative accessibility and reduced cellular heterogeneity as compared to other CNS regions. The major cell types of the retina are well-characterized and show a stereotyped and partially overlapping birth order, with retinal ganglion cells born first and Müller glia born last (Figure 1A). Both classic studies using retroviral-mediated sparse labeling of retinal progenitors in rodents (Turner and Cepko, 1987; Turner *et al.*, 1990), and more recent studies in both zebrafish and rodents using live imaging (Gomes *et al.*, 2010; He *et al.*, 2012) have shown that retinal progenitor cells (RPCs) are multipotent with changes in clone composition and size controlled largely by intrinsic mechanisms. These observations have raised several questions that remain unanswered (Cepko, 2014). First, how heterogeneous are RPCs, and do they exhibit any pre-specified cell fate biases? Second, how are changes in RPC competence controlled, and are competence transitions gradual or discrete? Third, what are the genes that direct an individual progenitor to exit the cell cycle and differentiate as a specific subtype of neuron or glia?

**Figure 1.**
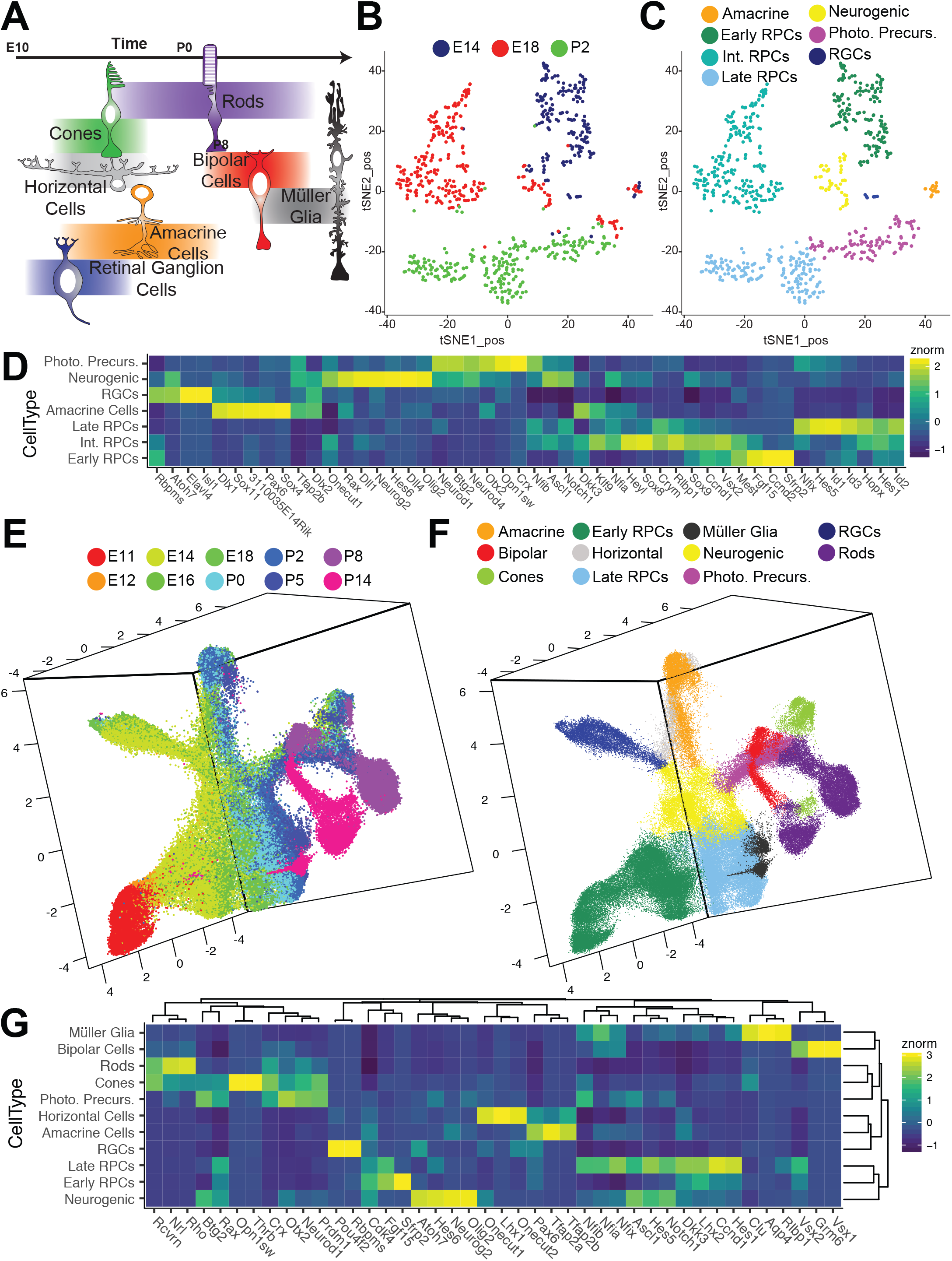
Single cell RNA-sequencing of the developing retina. (A) Schematic diagram of the temporal birth order of retinal cells. (B-C) tSNE-d¡mens¡on reduction of gene expression profiles from single Chx10:GFP-positive retinal progenitor cells via a modified Smart-Seq2 protocol with cells labeled by (B) developmental age or (C) annotated cell type as determined by marker gene expression in clustered cells. (D) Heatmap of select marker genes identified through differential expression analysis with respect to cell type on Smart-Seq2 samples. (E-F) UMAP-dimension reduction of droplet-based single cell RNA-sequencing of single developing mouse retinal cells with identified doublets and extra-retinal cells removed. Samples are colored by (E) developmental age or (F) annotated cell type as determined by marker gene expression in clustered cells. (G) Heatmap of differentially expressed genes across annotated developing retinal cell types in 10x samples. Abbreviations: Int. RPCs - Intermediate RPCs; Photo. Precurs. – Photoreceptor Precursors; RGCs – Retinal Ganglion Cells.

Past work provides some insights into each of these questions. Differential expression analysis has identified genes with dynamic expression amongst RPCs, both during the course of neurogenesis and at specific developmental stages (Blackshaw *et al.*, 2004; Trimarchi *et al.*, 2008). With few exceptions, the functional significance of this heterogeneity remains unresolved and confounded by bulk analysis of this dynamic cellular population. Cell lineage analysis of RPCs expressing individual transcription factors has revealed limited, and often temporally-regulated, cellular fate restriction (Brzezinski *et al.*, 2011; Brzezinski *et al.*, 2012; Emerson *et al.*, 2013; Hafler *et al.*, 2012; Jusuf *et al.*, 2012), but the relationship between RPC heterogeneity and cell fate determination remains unclear. Furthermore, although a limited number of transcription factors have been found to regulate RPC competence (Brzezinski *et al.*, 2012; Elliott *et al.*, 2008; Mattar *et al.*, 2015), these effects are either quite modest, or affect only retinal ganglion cell specification. Moreover, relatively little is known about the transcriptional regulatory networks that control the onset of retinal neurogenesis and the changes in mode of neurogenic division.

To comprehensively define the transcriptional changes and variation in gene expression associated with competence transitions, regulation of neurogenic divisions, and temporal cell fate specification, we conducted an extensive RNA-Seq analysis of the developing mouse retina at single cell resolution. We isolated single retinal cells at select time points ranging from prior to initiation of neurogenesis, through to its completion, including cells committed to each major cell type. Previous large-scale expression profiling studies of retinal development have been limited in their scope owing to the use of bulk dissected material (Aldiri *et al.*, 2017; Blackshaw *et al.*, 2001; Blackshaw *et al.*, 2004; Hoshino *et al.*, 2017). Past single cell expression profiling studies in retina either analyzed only small numbers of cells (Laboissonniere *et al.*, 2017; Mullally *et al.*, 2016; Trimarchi *et al.*, 2008), or profiled only mature cells in adult mice after fate specification (Macosko *et al.*, 2015; Shekhar *et al.*, 2016).

In this study, we examine the dynamics of gene expression within retinal cells in order to investigate the specifics of RPC competence, neurogenesis, and temporal cell fate specification. Using single cell RNA-sequencing (scRNA-seq), we are able to reconstruct developmental trajectories of distinct retinal cell fates, identify patterns of co-regulated genes involved in each of these developmental processes, and identify the genes and gene networks that directly influence RPC competence, retinal neurogenesis, and cell fate specification during development. We use these data to identify a comprehensive set of candidate genes that regulate progenitor competence and retinal neurogenesis, and we identify the NFI family of transcription factors as essential for generation of late-born retinal cells as well as termination of RPC proliferation. This work advances our understanding of retinal development, and additionally provides a template for investigation of temporal patterning in all areas of the developing CNS.

## RESULTS

### Examination of Retinal Progenitor Cell Heterogeneity via Smart-Seq Analysis

To obtain a detailed transcriptional profile of single RPCs over the course of retinal neurogenesis, we first performed single cell RNA-sequencing on FACS-isolated Chx10-GFP+ mouse RPCs (Rowan and Cepko, 2004), using an adapted Smart-Seq2 protocol (Chevee *et al.*,2018) at embryonic (E) days 14 and 18, and postnatal (P) day 2, which correspond to early, intermediate and late stages of retinal neurogenesis, respectively (Figure 1B). Analysis of 780 individual RPCs profiled to a depth of ~750,000 aligned fragments per cell (mean 762,726; sd 291,612.5; Figure S1A-D) revealed that these cells resolved into three major clusters that express canonical RPC markers (e.g. Ccnd1, Cdk4, Pax6; Figure S1F), and correspond to each of the time points sampled (Figure 1B; Figure S1E) when plotted using 2-D t-stochastic neighbor embedding (tSNE) analysis (van der Maaten and Hinton, 2008) using genes that displayed high-variance in expression across all cells (Table S1). We further resolved these clusters into discrete groups distinguished by expression of genes specific to distinct phases of the cell cycle, with subclusters specific to G1/S (e.g. *Pcna,Ccne2*) and G2/M phase (e.g. *Ccnb1, Ube2c*), respectively (Figure S1G). A much smaller and distinct cluster, which included cells from each age, expressed both genes associated with active proliferation *(Dkk3, Cdk4)* and multiple neurogenic bHLH factors (e.g. *Atoh7, Olig2, Neurog2, Neurod1;* Figure S1H). Finally, in the P2 sample, a substantial subset of cells lacked expression of genes associated with proliferation, and instead expressed markers of immature photoreceptors (e.g. *Crx*), while other small clusters of postmitotic cells expressed genes specific to immature amacrine (*Tfap2b*) and retinal ganglion cells (*Pou4f2, Isl1;* Figure S1I). Therefore, we annotated the cell type of each individual cell based on transcriptional profiles of individual clusters (Figure 1C).

Differential gene expression testing across the non-neurogenic progenitors identifies 1195 genes (q-val < 1.0 e-10, mean expression > 1.0; Table S2) with significant differential expression amongst progenitor populations across development. This suggests that RPCs show substantial and global changes in gene expression over the course of development. However, within individual ages, we do not observe subclusters amongst these RPCs, except for expression of transcripts associated with cell cycle phase. Conversely, differential expression testing across annotated cell types identified 4754 genes (q-val < 1.0 e-10; Table S3) including several genes known to promote retinal neurogenesis and photoreceptor specification (Figure 1D). Genetic lineage analysis has suggested that RPCs expressing neurogenic bHLH factors are more likely to undergo terminal neurogenic divisions (Brzezinski *et al.*, 2011; Brzezinski *et al.*, 2012; Hafler *et al.*, 2012). Together, these results indicate that RPCs undergo significant transcriptional changes across developmental time, consistent with a change in developmental competence, and that both cell cycle phase and neurogenic potential significantly influence the transcriptional heterogeneity of RPCs at any given time point.

### Droplet-based scRNA-Seq reveals the full transcriptional landscape of mouse retinal development

To build upon these initial findings, we next transitioned to droplet-based single cell RNA sequencing to increase the cellular resolution across retinal development by expanding both the number of cells and time points. We profiled 120,804 single cells from whole retinas at 10 select developmental time points, from retinal neuroepithelial stages (E11) through terminal fate specification (P14), using the 10x Genomics Chromium 3’ v2 platform (PN-120223) (Figure S2A). Libraries were pooled and sequenced on the NextSeq 500 platform to a mean depth of ~110,220,000 reads per library, corresponding to a mean UMI count of 2099.75 and 1153.43 genes per cell (Figure S2B-E). Raw data were processed through the Cell Ranger pipeline and raw counts were aggregated together for input into the Monocle R/Bioconductor platform (Trapnell *et al.*, 2014). Preliminary clustering and cell type annotation was performed on single cell profiles obtained from individual timepoints using a modified Monocle dpFeature workflow (Qiu *et al.*, 2017) (Figure S3-S4). All time points were then aggregated into a single cell dataset for further analyses. Using a set of 3290 high-variance genes across all cells (Table S4), we established a reduced three-dimensional representation of the developing retina using UMAP (Mclnnes and Healy, 2018) (Figure S2F-G; Movie 1-2). A second round of clustering (Figure S2H) and cell type annotation was performed at which time obvious clusters of doublets and extra-retinal cells were identified and removed (Figure 1E-F; Figure S2I; Movie 3).

The resulting representation contains a core manifold consisting of retinal progenitor cells at all ages between E11 and P8 that express canonical RPC markers *(Pax6, Vsx2, Lhx2*, etc; Figure 1G). We also observe a population of mitotically active *(Ccndl* and Dkk3-positive) cells that express multiple neurogenic bHLH genes *(Olig2, Neurog2)*with reduced expression of *Vsx2* and *Lhx2* compared to other RPCs (Figure 1G). This population corresponds to the neurogenic RPC population identified in the Smart-Seq analysis (Figure 1C-D) and is observed exclusively between E12 and P8. The neurogenic population is immediately adjacent to, and extends from, the main RPC group (Figure 1E-F). Trajectories of differentiating cells corresponding to all major retinal neuronal subtypes, with the exception of horizontal cells, can be seen directly emerging as separate branches from this population of neurogenic RPCs. A branch corresponding to differentiating Müller glial precursors, in contrast, emerges directly from the main cluster of RPCs and not through the neurogenic phase, in line with Müller glia and RPCs exhibiting largely overlapping gene expression profiles (Blackshaw *et al.*, 2004; Nelson *et al.*, 2011; Roesch *et al.*, 2008). Examination of gene expression within this representation identified cells with high levels of horizontal cell-specific gene expression *(Lhx1* and *Lhxlos)* present within the presumptive amacrine trajectory. Recursive analysis of the amacrine trajectory (subsetting, identification of high variance genes, dimension reduction and sub-clustering) resulted in identification of both starburst amacrine and horizontal cells as separate populations distinct from other developing amacrine cells (Figure S5A-B).

Within the reduced dimensional embedding, the great majority of cells are distributed along a single contiguous manifold, indicating that we have profiled the majority of key transitions that occur during the course of mouse retinal development. Cell-type classification, number and proportions of annotated cell types are listed in Table S5 and Figure S2J-L. The observed timing and sequential progression of major cell type trajectories is also consistent with previous studies (Blackshaw *et al.*, 2004). Examinations of the proportions of RPCs, neurogenic progenitors and gliogenic cells revealed a relatively stable proportion of neurogenic cells captured during retinal neurogenesis from E14-P8 (Figure S2M). After P2 the fraction of gliogenic cells increases, consistent with an increase in terminal divisions as neurogenesis progresses, as observed in time-lapse imaging in previous studies (Gomes *et al.*, 2010; He *et al.*,2012).

To identify the set of genes with dynamic regulation during the course of retinal development, we performed a pseudotemporal analysis using 3259 high variance genes in neuroretinal cells only, excluding both annotated extraretinal cells and doublets (Trapnell *et al.*, 2014) (Table S6). Due to matrix size limitations of dependent algorithms, we performed the pseudotime analysis on ~32,000 cells randomly sampled across the entire dataset. Using this approach, we were able to reconstruct a complex tree that largely reflects known temporal ordering of cell fate specification within the retina, and displays terminal branches of the tree that are both predominantly comprised of single cell types and reflect the known developmental trajectories of these cell (Figure 2A-B; S6).

**Figure 2.**
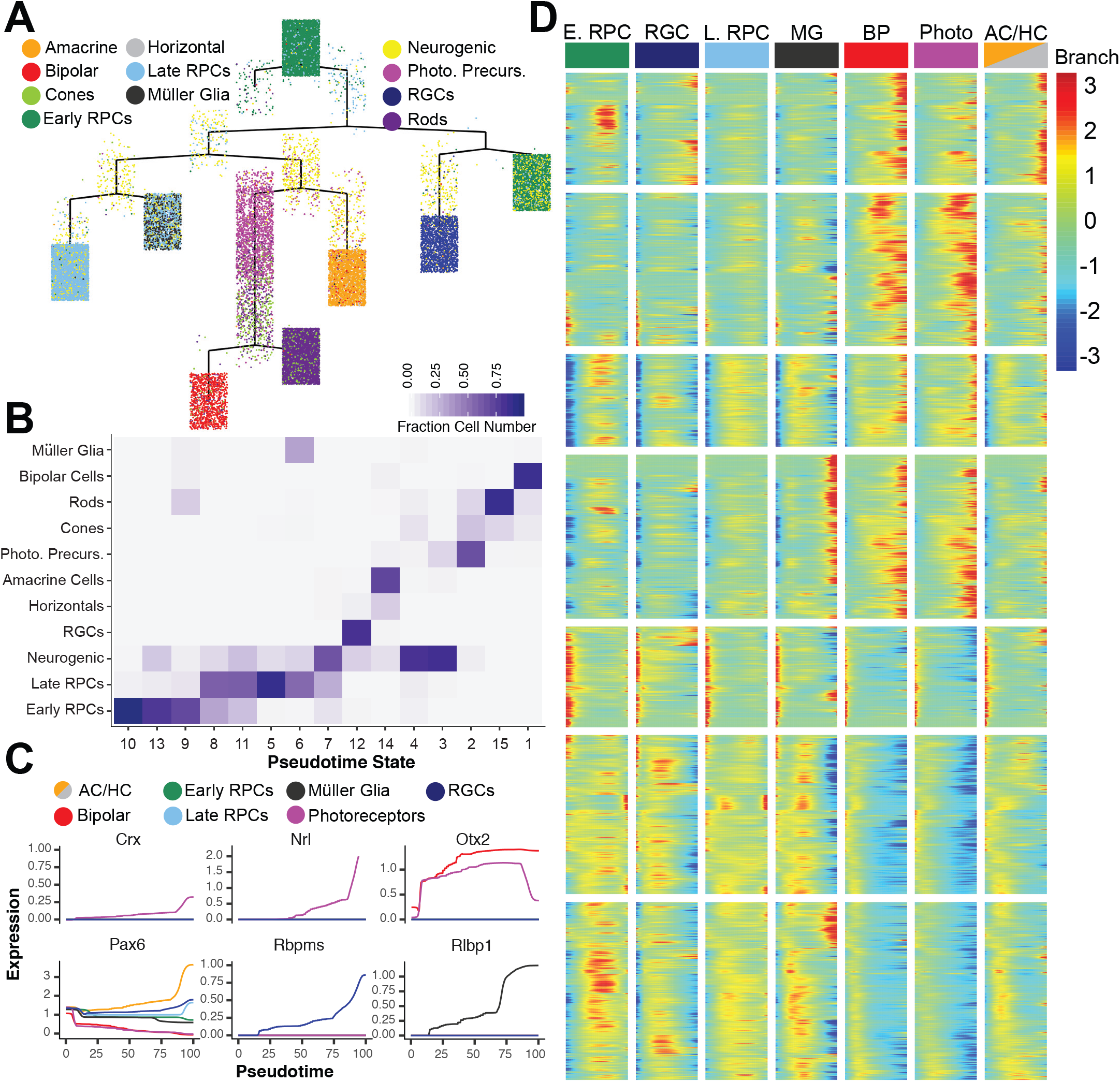
Pseudotime analysis reveals temporal regulation of transcript expression during differentiation of retinal cell types. (A) Complex pseudotime tree on a subset of retinal cells colored by annotated celltype. (B) Heatmap of pseudotemporal states reveals the proportion of individual cell types assigned to each state. (C) Known marker gene expression across pseudotime colored by cell type designation of each trajectory. (D) Branched heatmap faceted by terminal pseudotime state corresponding to annotated cell type. Expression of all 7,487 differentially expressed transcripts across pseudotime starting from pseudotime value of 0 (left side of each column; top of the complex tree in panel A), and following a continuous path down the tree towards each terminal branch (right side of each column. Abbreviations: AC/HC - Amacrine Cells/Horizontal Cells; BP - Bipolar Cells; E. RPC - Early RPC; L. RPC - Late RPC; MG - Müller Glia; Photo - Photoreceptors; Photo. Precurs. - Photoreceptor Precursors; RGCs - Retinal Ganglion Cells.

Differential gene expression analysis across pseudotime identified a total of 7487 genes with significant changes in gene expression over the course of retinal development (Figure 2D; Table S7 q-val < 1e-5). This list includes known markers of maturing cell types (Figure 2C). Additionally, recursive analysis of major branch points in the complex pseudotime tree identified genes that exhibit significant differential expression during the specification and maturation of individual cell fates. Using this strategy, we were able to identify genes that showed selective expression during early, intermediate and late stages of differentiation of postmitotic precursors of retinal ganglion cells, amacrine/ horizontal cells, photoreceptor cells, bipolar cells and Müller glia (Figure S7-S12). Temporal- and trajectory-appropriate expression of all transcription factors known to regulate differentiation of individual retinal cell types was observed, with *Isl1* and *Pou4f2* readily detected as early markers of RGCs (Li *et al.*, 2014), and *Otx2* for immature photoreceptors and bipolar cells (Baas *et al.*, 2000), for instance. Additional recursive analysis of the amacrine/horizontal cell trajectory further identified genes that selectively mark developing amacrine, starburst amacrine, and horizontal cells (Figure S5C-D; Figure S13-14).

### Temporally dynamic changes in gene expression within RPCs

To characterize the fundamental transcriptional changes within RPCs across developmental time, as well as the changes in gene expression within progenitors likely to commit to the neurogenic or gliogenic fractions of cells, we performed a pseudotemporal analysis on subsets of specific cell types. We first addressed the changes in gene expression across development in the RPC subset. UMAP representations and pseudotime analyses using a set of 1763 high variance genes across the subsetted RPCs revealed a temporal progression from early (E11) through late (P8) time-points. Interestingly, we observed a clear segregation of RPCs that occurs between E16 and E18. A clustering solution for the cells in UMAP space agrees with this classification and allowed us to delineate stable classes of early and late RPCs (Figure 3A-C; Figure 1F). Importantly, cells from both E16 and E18 timepoints were stratified across this divide, indicating that this difference likely does not reflect a significant batch effect, but rather a major physiological transition within the RPCs themself (Figure 3C). The stratification of early vs late RPCs is consistent with a developmental window (E16-E18) that is well correlated with the end of a developmental competence phase in which early-born retinal cell types (RGCs, horizontal cells, cone photoreceptors and GABAergic amacrine cells) are generated (Voinescu *et al.*, 2009; Young, 1985a, b)(Figure 1A). We identified 3291 genes that display significant differential expression across the RPC pseudotime (qval <1e-20, Figure 3E). Notably, we identify previously established markers of both early (*Fgf15* and *Sfrp2*) and late (*Crym* and *Car2*) RPCs (Blackshaw *et al.*, 2004) and a host of previously unidentified markers of early versus late RPCs, including the NFI family transcription factors (*Nfia, Nfib*, and *Nfix;* Figure 3E).

**Figure 3.**
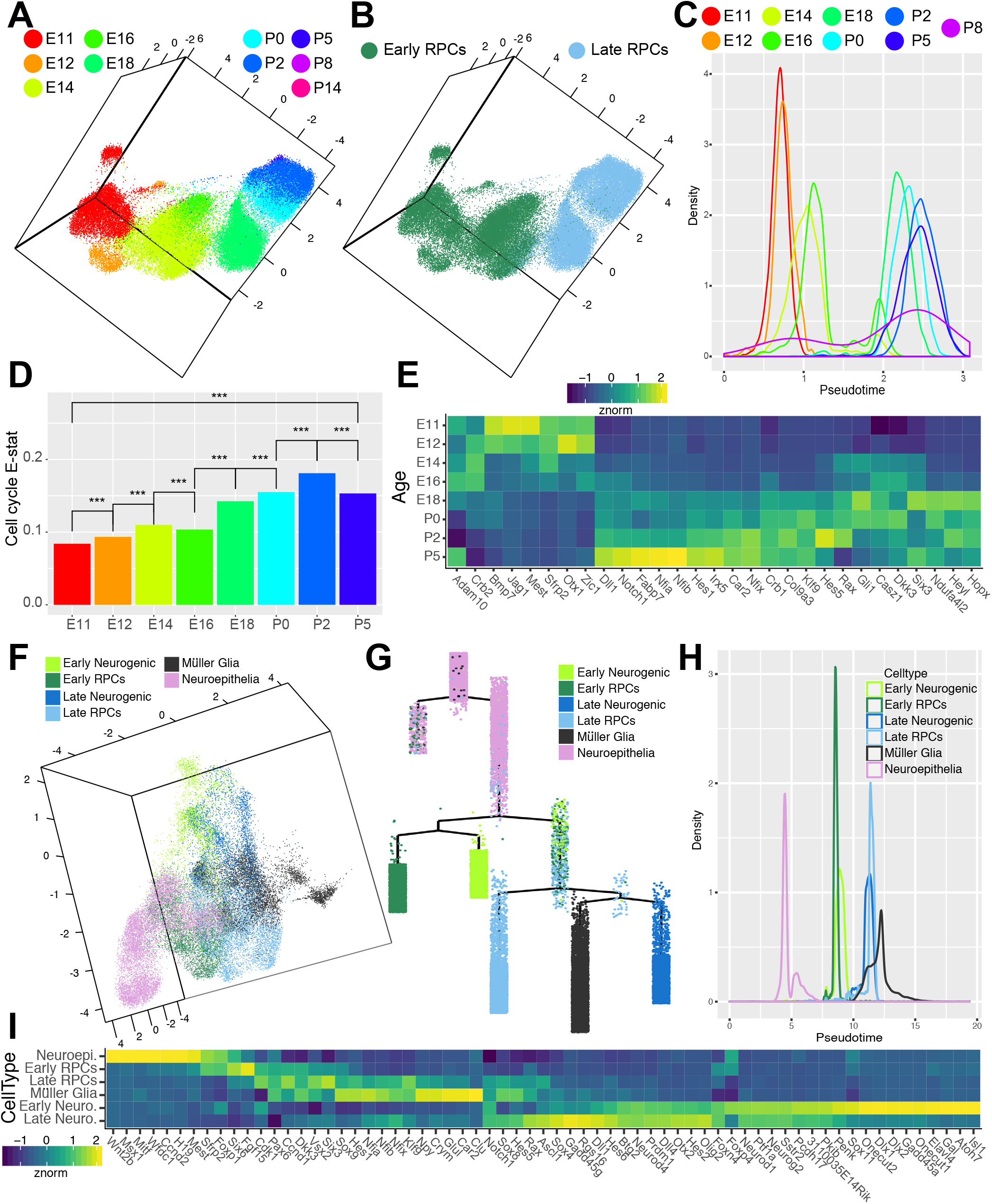
Identification of candidate transcripts in the regulation of RPC competence, neurogenesis and gliogenesis. (A-B) UMAP dimension reduction of subsetted RPCs colored by (A) developmental age or (B) annotated celltype. (C) Density plot of cells across RPC pseudotime analysis and colored by developmental age. (D) EVA analysis of relative dissimilarity of canonical cell cycle genes within RPCs assessed at individual developmental ages. (E) Heatmap of differentially expressed genes across RPCs pseudotime, displaying normalized transcript enrichment across cells grouped by developmental age. (F) UMAP-dimension reduction displaying the classification of neuroepithelial cells, early and late RPCs, early and late neurogenic, and gliogenic cells identified in pseudotemporal analyses. (G) Complex pseudotime tree of RPCs, neurogenic and gliogenic cells, highlighting designation of neuroepithelial cells, early and late RPCs, early and late neurogenic cells, and Müller glia. (H) Density plot of RPC, neurogenic, gliogenic, and neuroepithelial cells across pseudotime. (I) Heatmap of differentially expressed genes across RPC/neurogenic/gliogenic pseudotime, displaying normalized enrichment across all cells within the designated celltype. *** indicates p<0.001 from EVA analysis.

To further examine the heterogeneity of the RPC population across development, we applied Expression Variation Analysis (EVA) to quantify the relative dissimilarity in transcriptional profiles between RPCs from distinct developmental time points (Afsari *et al.*, 2014). Briefly, EVA provides a robust statistical framework, quantifying differences in heterogeneity of expression within pathways between experimental conditions, that is directly applicable to imputed scRNA-seq data. We observe that variation in expression of cell cycle genes (Figure 3D) and the FGF pathway (Figure S15C) increases over developmental time, whereas variation in the Wnt and Notch pathways decreases over time (Figure S15C). This observation is consistent with dysregulation in cell cycle and FGF pathways over developmental time, and it is also consistent with regulation of Wnt and Notch pathways at later stages. Of particular note, we suggest that the increased variance in cell cycle gene expression in late-stage RPCs likely serves as a proxy signal for the increase in cell cycle length in RPCs that is seen over the course of development, as previously observed in the rat retina (Alexiades and Cepko, 1996).

Strikingly, Notch pathway gene expression increased steadily in RPCs over the course of retinal development, forming a smooth temporal gradient that peaked in late-stage RPCs. This is particularly clear when canonical Notch target genes such as *Hes1* and *Hes5* are examined (Figure 3E; Figure S15A), and is consistent with the known role of Notch signaling in driving specification of late-born Müller glia (de Melo *et al.*, 2016a; de Melo *et al.*, 2016b; Jadhav *et al.*, 2006). The temporal regulation of Notch pathway gene expression observed within the dataset is consistent with the EVA analyses, which suggests a more homogeneous activation of the pathway within RPCs as development progresses (Figure S15C).

We also observed a smaller number of genes that show altered expression in RPCs from P0 onwards. Downregulated genes include transcription factors and chromatin-associated proteins (*Baspl*), as well as known cell cycle regulators (*Kpna2, Kif2a*), while a limited number of Muller glia-expressed genes are upregulated *(Cd9, Sat1;* Figure S15B). Interestingly, a number of genes also show elevated expression in early (E11-E12) RPCs and postnatal RPCs (*Oaz1, Pebpl;* Figure S15B). The functional significance of these changes may reflect the increased length of the cell cycle that is seen in late-stage RPCs, or simply the loss of proliferative potential of late RPCs, as the vast majority of RPCs are undergoing terminal mitoses (Alexiades and Cepko, 1996; Turner and Cepko, 1987).

Since we had observed significant changes in gene expression across pseudotime in RPCs, we next investigated the potential of these gene expression differences to translate into functional differences likely to regulate neurogenesis or gliogenesis. Therefore, we subset the data to all RPCs (both neurogenic and non-neurogenic) or gliogenic cells and again performed pseudotemporal analysis on ~32,000 sampled cells. Analysis of the resulting complex pseudotemporal hierarchy confirmed separate early and late progenitor populations, as well as the gliogenic population. However, three additional populations of cells were identified based on differential expression of genes across pseudotime states. A neuroepithelial/ciliary margin population emerged, comprising cells primarily from the earliest (E11-E14) time points and prior to the onset of RPC neurogenesis (Figure 3F-H; Figure S15D-F). Genes with enriched expression within this subset of cells include those associated within ciliary margin (*Msx1, H19*) and/or retinal pigmented epithelium (*Mitf, Ccnd2*), and reflect genes expressed after early eye-field specification (Blackshaw *et al.*, 2004)(Figure 3H).

The two additional populations of cells observed were a bifurcation of the neurogenic population, corresponding to early and late neurogenic subtypes. The subdivision of the RPC population into early and late RPCs around E16-E18 was recapitulated in an early and late population of neurogenic cells. Several genes are selectively expressed in either early (*Gadd45a, Sox11, Elavl3, Gal*) or late-stage (*Gadd45g, Rgs16*) neurogenic cells (Figure 3I). Notably, transcription factors previously reported to control specification of early-born cell types such as RGCs (*Sox11, Atoh7*), horizontal cells (*Onecut1/2*) and GABAergic amacrine cells (*Dlx1/2*) were selectively expressed in early neurogenic RPCs (de Melo *et al.*,2005; Emerson *et al.*, 2013; Jiang *et al.*, 2013)(Figure 3I). Likewise, genes associated with specification of late-born cell types (*Prdml, Otx2, Ascll*) were enriched in the late-neurogenic fraction (Brzezinski *et al.*, 2010; Katoh *et al.*, 2010; Nelson *et al.*, 2009; Nishida *et al.*, 2003) (Figure 3I). These transcriptional differences suggest a sharp functional distinction between early and late neurogenic cells that derives in part from transcriptional differences established in early vs late retinal progenitors, and is consistent with a change in progenitor competence over the course of retinal development.

### Identification of co-regulated patterns of gene expression across retinal development

While our supervised pseudotemporal analyses identified developmental trajectories associated with most major retinal cell types, it was unable to resolve more closely-related cell types or trajectories with a high degree of gene reuse; e.g. when the same gene is used in multiple different pathways and/or cells, such as the differentiation of horizontal cells from amacrine cells and the differences between immature cone and rod photoreceptors. Gene reuse poses an even greater analytical challenge in development where marker genes can implicate different biological processes depending on timing and context (Stein-O’Brien *et al.*, 2017a). Previously, an unsupervised smooth-sparse Bayesian non-negative matrix factorization (NMF) algorithm, CoGAPS, has successfully parsed gene signatures of discrete pathway use and highly related cell types in bulk RNA-Seq despite highly overlapping gene networks (Fertig *et al.*, 2010; Stein-O’Brien *et al.*, 2017b). To account for gene reuse in our single cell dataset, further validate our supervised analysis and identify gene signatures of discrete cell populations, we expanded and reengineered CoGAPS for scRNA-seq data in a new algorithm called scCoGAPS(Stein-O’Brien *et al.*, in prep). Unlike other methods which learn gene signatures on a subset of cells, and then project the entire dataset (Macosko *et al.*, 2015), scCoGAPS uses an ensemble-based approach across multiple sets of cells to learn these signatures in parallel across 120,804 cells. This ensemble-based approach has the additional advantage of providing measures of signature robustness and significantly increased computational speed. Furthermore, Bayesian NMF algorithms have previously been shown to more accurately parse the cumulative effect of genes that participate in multiple pathways by directly encoding the dependence of biological processes (Kossenkov *et al.*, 2007; Ochs and Fertig, 2012). The sparsity constraint inherent in the CoGAPS engine of scCoGAPS further ensures robustness by accounting for biological parsimony, and was developed to specifically account for the additional sparsity and distribution of scRNA-Seq data.

scCoGAPS decomposes the input data into two matrices: a pattern matrix with relative cells weights along rows and an amplitude matrix with relative gene weights along columns. Each row of the pattern matrix quantifies the extent of transcriptional changes within the genes in the corresponding column of the amplitude matrix. The columns of the amplitude matrix and rows of the pattern matrix provide a low dimensional representation of the biological processes, or latent space, in that data. Gene reuse is captured by allowing genes to have large amplitude values in multiple columns, and thus associated with multiple biological processes or states. Identification of genes uniquely associated with a given latent space can be performed with our PatternMarker statistic (Stein-O’Brien *et al.*, 2018; Stein-O’Brien *et al.*, 2017b). Here, we apply the scCoGAPS algorithm and the PatternMarker statistic to associate patterns with known biological processes in scRNA-seq data and use this for novel biomarker discovery.

We identified a total of 80 scCoGAPS patterns across the full developmental time course, including patterns that selectively labeled differentiating horizontal cells from amacrine cells as well as cones from rods, and identified several co-regulated gene networks that reflect cell type specification, key developmental transitions, and correlate with phenotypic features of the dataset (Figure 4A; Figure S16). PatternMarker genes and corresponding gene weights for each of the identified 80 patterns are indicated in Tables S8-9. Importantly, patterns with high weights in annotated horizontal cells (Patterns 2 + 16) correlated well with our manual annotation, and also highlight gene reuse across specification of disparate cell types, exhibiting high pattern weights in a subset of mature RGCs (Figure S16). Additional patterns, for example, are associated with mature RGCs (Pattern 15; Figure 4B), or recover other phenotypic features of these data such as sex (Pattern 36; Figure 4C, Movie 4).

**Figure 4.**
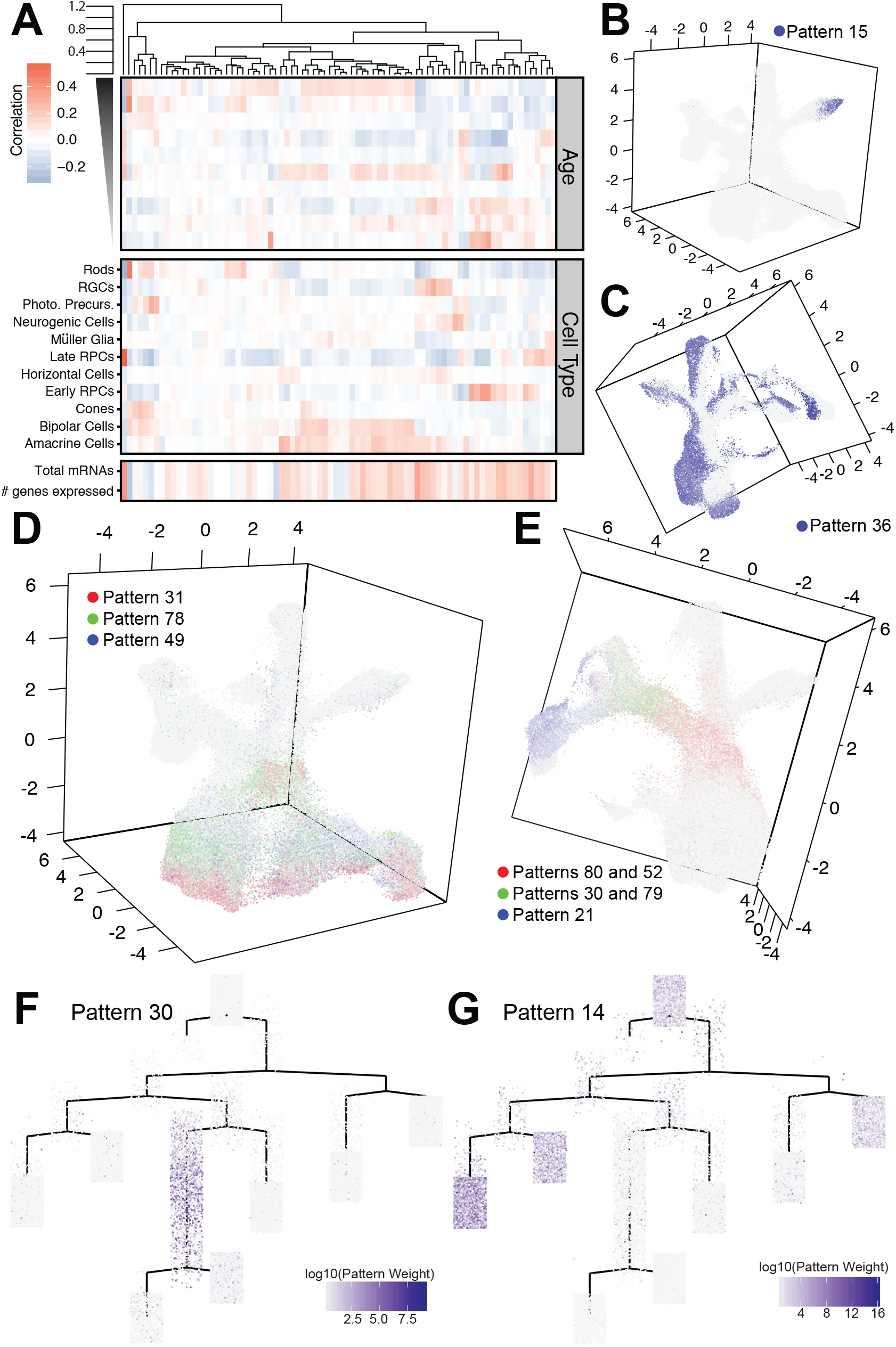
scCoGAPS analysis on single cell RNA-sequencing samples reveals patterns of gene expression within developmental processes. (A) Correlation heatmap of pattern weight with annotated phenotypic data within the dataset including biological parameters (cell type annotation and developmental age) and technical artifacts (sequencing depth) and patterns relatedness and clustering indicated according to the dendrogram at the top. (B-E) Examples of graphical representations of pattern weights of individual cells within the UMAP-dimension reduction. (B) Pattern 15 marks the terminal trajectory of RGCs. (C) Pattern 36, with pattern marker Xist, highlights the sex of the animals that cells came from. (D-E) Combinations of patterns can be used to assess (D) the influence of cell cycle phase on clustering of RPCs or (E) developmental transitions across clusters, from neurogenic cells through photoreceptor specification and differentiation. (F-G) scCoGAPs pattern weights of cells plotted within the retina complex pseudotime (Figure 2A) highlights (F) photoreceptor/bipolar cell precursors by Pattern 30’ using Otx2 as a pattern marker, and (G) late RPCs cells with high pattern weights in Pattern 14.

Visualization of multiple, temporally-regulated patterns can be used to identify and characterize continuous biological processes such as the progression of progenitors through the cell cycle or the specification of individual cell fates over developmental time (Figure 4D-E; Movies 5-6). Plotting pattern weights on pseudotime representations (Figure S17) highlights the association of patterns with developmental transitions (Figure 4F; photoreceptor/bipolar cell precursors with high *Otx2* expression) or differences between RPC states (Figure 4G). The concordance of the data-driven patterns not only validates the supervised pseudotemporal analysis, but also provides a more focused and unbiased list of biomarkers. The application of scCoGAPS to single cell RNA-Seq data also captures technical aspects of these data as well. Combinations of biologically incompatible scCoGAPS patterns (e.g. two patterns for distinct mature cell types within the same cell) can readily delineate doublet cell populations. Thus, the ability to account for gene reuse makes scCoGAPS a powerful new method for data driven scRNAseq analysis.

### NFI family transcription factors both regulate specification of late-born retinal cell types and drive cell cycle exit

A consistent theme in our differential expression, pseudotime, and scCoGAPS analyses was the identification of the NFI family of genes, specifically *Nfia/b/x*, as having high expression and high amplitude weights in cells/patterns associated with late-stage RPCs, bipolar cells and Müller glia (Figure S18). We also observed expression of Nfia/b/x within the amacrine cell trajectory, consistent with previous reports of *Nfia* expression within aII amacrine cells (Keeley and Reese, 2018; Figure S18). This expression pattern was confirmed via in situ hybridization (Figure 5A). *Nfia* has been previously found to be necessary for astrogliogenesis in both the cortex and spinal cord (Deneen *et al.*, 2006; Kang *et al.*, 2012; Nagao *et al.*, 2016), and to promote the formation of cells expressing a subset of Müller glial markers (de Melo *et al.*, 2016a). We hypothesized that the NFI family of transcription factors may play a similar role in mediating the transition from neurogenic to gliogenic commitment in the developing retina. Based on these observations, we chose to further investigate the function of the NFI family of transcription factors within late-stage RPCs.

**Figure 5.**
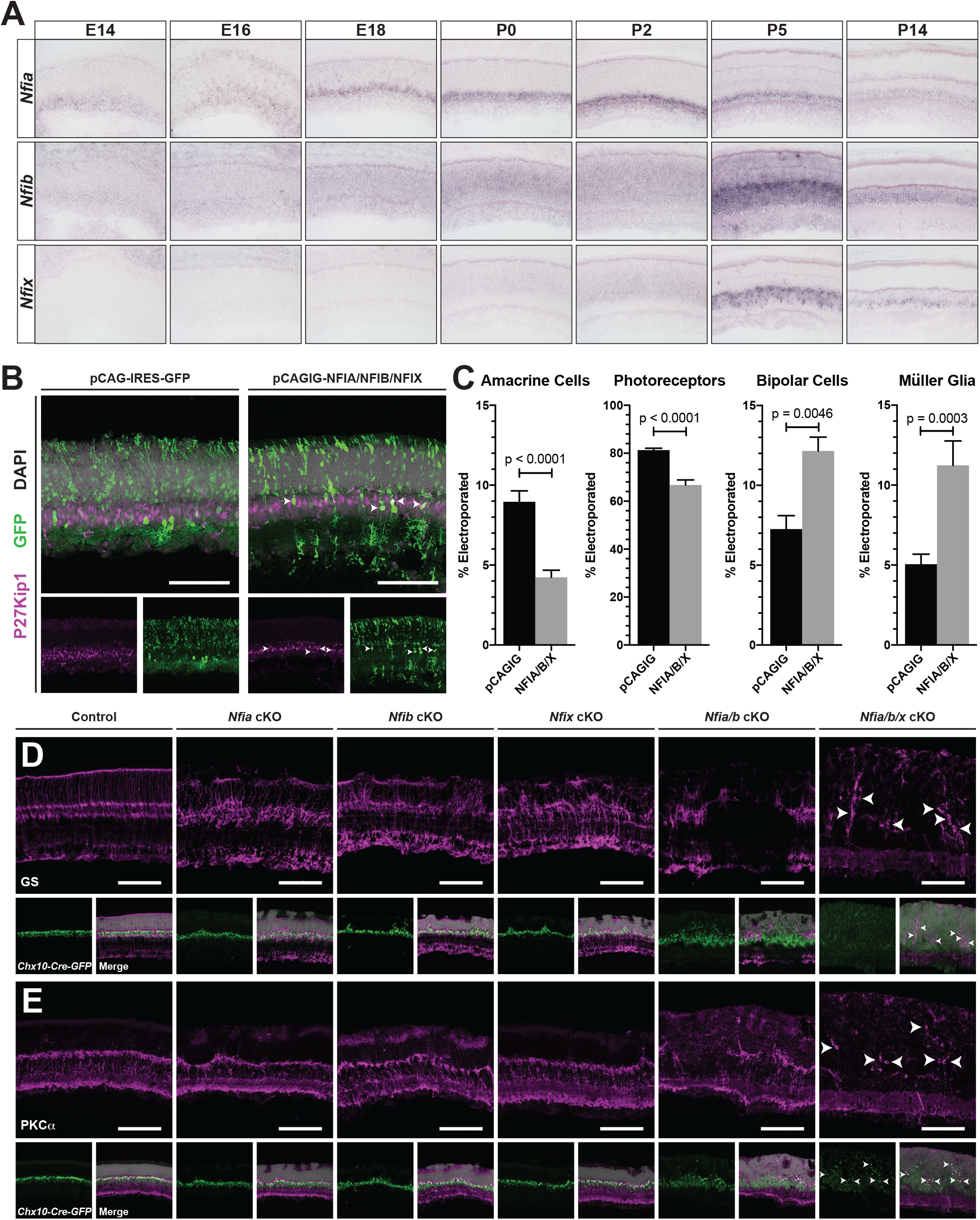
The NFI family of transcription factors regulate specification of late retinal cell fates. (A) Time-series of in situ hybridization expression analysis of *Nfia, Nfib*, and *Nfix.* (B) Examples of P14 retinas subjected to immunohistochemistry for the Müller glia marker P27Kip1 after in vivo electroporations at P0 of either control (pCAGIG) or the combination of pCAGIG-NFIA, NFIB, and NFIX constructs. (C) Quantification of the proportion of electroporated cells that co-label with markers for amacrine, photoreceptor, bipolar and Müller glial after in vivo electroporation of pCAGIG control or pCAGIG-NFIA/B/X. (D-E) Chx10-Cre mediated loss of function of *Nfia, Nfib, Nfix, Nfia* and *Nfib*, or *Nfia, Nfib, Nfix.* (D) Disruption of retinal organization and loss of Müller glia is marked by glutamine synthetase staining (GS). (E) Loss of bipolar cells is observed through PKCα staining, a marker or rod-bipolar cells. Arrowheads indicate remaining Müller glia (D) and bipolar cells (E). p-values are the result of unpaired t-tests on cell counts. Cell counts were performed on 2 or more sections from 5-8 animals per condition. Scale bars - 100μm

We used in vivo electroporation to overexpress NFIA/ B/X genes in P0.5 retina (de Melo and Blackshaw, 2018), and, at P14, observed an increase in both Müller glia and bipolar cells -- the two last-born major retinal cell types -- along with a complementary reduction in the fraction of rod photoreceptors and amacrine cells (Figure 5B-C). By mating conditional alleles of each NFI gene (Campbell *et al.*, 2008; Hsu *et al.*, 2011) to the RPC-specific *Chx10*-Cre line (Rowan and Cepko, 2004), we generated RPC-specific single, double and triple mutants of *Nfia, Nfib*, and *Nfix*. We observed that RPC-specific loss of function of either *Nfia, Nfib* or *Nfix* led to a disruption of Müller glial marker staining and inconsistencies within the retinal outer limiting membrane (OLM) (Figure 5D; Figure S20B). These phenotypes were amplified in *Nfia/b* double mutant mice, exhibiting a more pronounced loss of Müller glial markers (glutamine synthetase; GS; Figure 5D; p27, Lhx2; Figure S19) and a severe disruption of the OLM, which led to major defects in retinal lamination (Figure S20B). These animals also showed a substantial reduction in bipolar cell markers including Pkcα, Vsx2, and Isl1 (Figure 5E; Figure S19). *Nfia/b/x* triple mutants showed a nearly complete loss of both Müller glial and bipolar cell markers (Figure 5D-E), although other retinal cell-specific markers were preserved (Figure S19). Taken together, these results demonstrate that *Nfia/b/x* are necessary and sufficient for specification and differentiation of late-born retinal cell types, the Müller glia and bipolar cells.

In addition to the significant loss of bipolar and Müller glial cells, we observed a maintenance of weak *Chx10*-GFP-Cre reporter expression within both Nfia/b double and *Nfia/b/x* triple mutant retinas. This weak transgene expression, compared to the strong bipolar cell transgene expression, is similar to the expression levels of the transgene observed in RPCs. Based on the Chx10-GFP-Cre transgene expression and previous reports implicating Nfix in proliferative quescience (Martynoga *et al.*, 2013), we hypothesized that the NFI mutants may retain a population of proliferating RPCs. To address this question, we analyzed the RPC marker Mki67 within the mutant retinas. We noted that proliferation persisted in the retinas of P14 *Nfia/b* double mutants and *Nfia/b/x* triple mutants, well beyond ages when RPCs are normally present (Figure 6A-B). Proliferating cells were abundant even as late as P28 (Figure 6C, Figure S20F), accompanied by thickening of the retina (Figure S21D). Although a small number of proliferating microglia are observed in these mutant retinas (Figure S20C+E), we observe co-labeling of most proliferative cells with the Chx10-GFP-Cre transgene with Mki67, suggesting that the proliferative cells are of retinal origin (Figure 6C). However, these cells are practically devoid of other canonical RPC markers such as Pax6 and Vsx2, and also fail to express RPC-specific genes that are also required for Müller glial development such as Lhx2 (de Melo *et al.*, 2016b; Figure S19). Consistent with the activation of microglia within the retina, remaining Müller glial cells within the retinas of *Nfia/b*double or *Nfia/b/x* triple mutants have become reactive, exhibiting increased expression of glial fibrillary activating protein (GFAP; Figure S20A).

**Figure 6.**
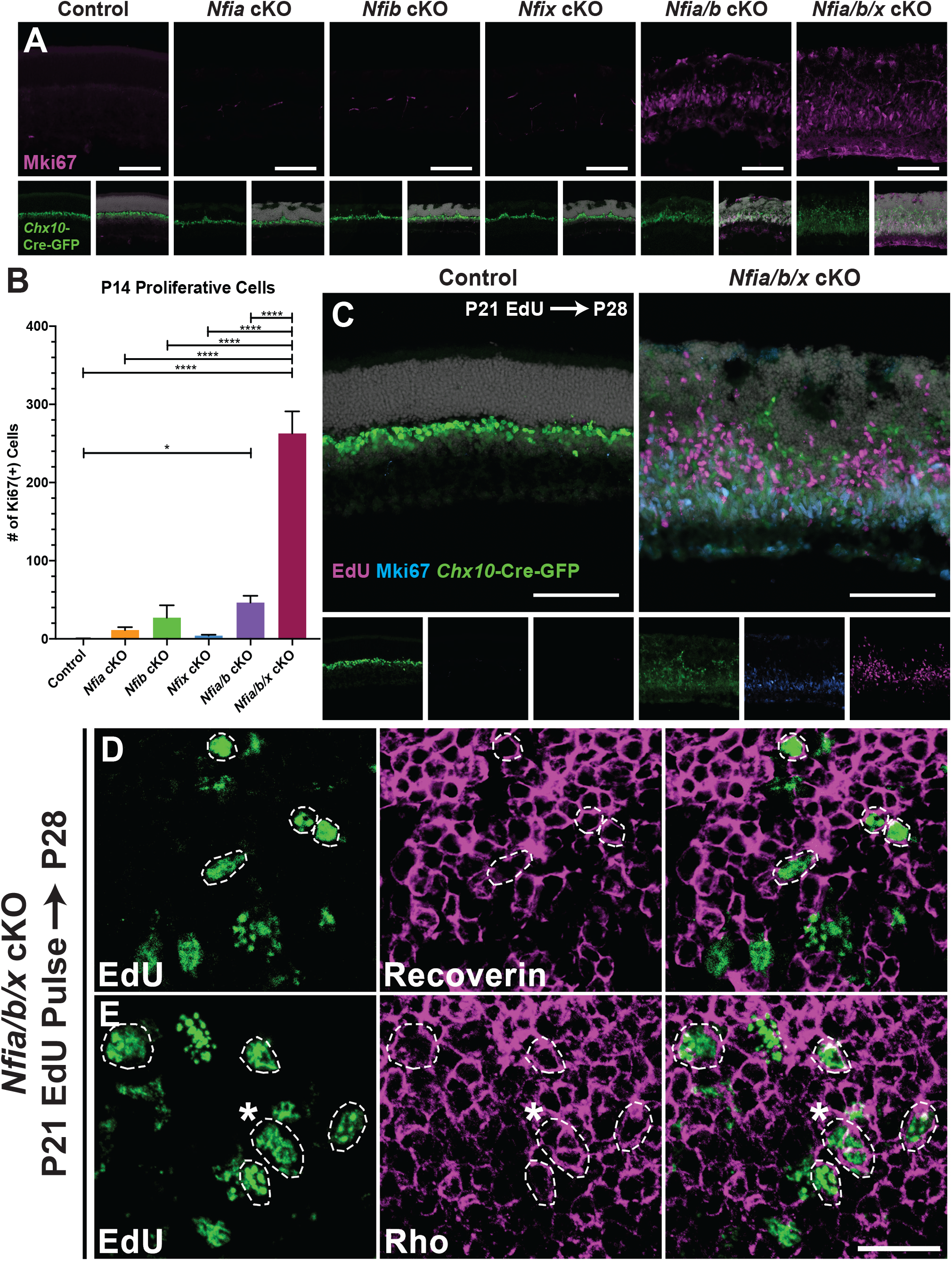
Loss of NFI transcription factors results in sustained RPC proliferation and neurogenesis. (A) Proliferative cells were observed through staining of retinas with MKi67 across genotypes. (B) Quantification of Ki67 cells; p < 0.0001 (C) P21 EdU injections chased to P28 and co-stained with Ki67 in control and *Nfia/b/x* triple mutants. (D-E) Confocal images of retinas from P21 EdU injections at P28 and co-labeling with (D) recoverin or (E) rhodopsin indicates differentiation of ectopic RPCs into photoreceptors (circled nuclei). Asterix in panel E indicates 3 nuclei that display colocalization. Statistics are results of a one-way ANOVA followed by Tukey’s multiple comparison test. * - p < 0.05; **** - p < 0.0001. Scale bars: A and C - 100μm; D-E - 20μm

To examine the extent of differentiation of this ectopic population of proliferative cells within the retina, we pulsed control and *Nfia/b/x* triple mutants with EdU at P21 and chased labeled cells at P28. EdU labeling indicates that ~15% of proliferating cells in P21 retina give rise to recoverin-positive photoreceptor-like cells by P28 (Figure 6D-E; Figure S20G). However, a proportion (~7%) of EdU labeled cells maintain expression of the proliferative marker Mki67 (Figure 6C; Figure S20G). Therefore, retinal thickening observed in *Nfia/b/x* triple mutants can, at least in part, be attributed to the continuous proliferation and differentiation of retinal cells. Together, this indicates that expression of *Nfia/b/x* in late-stage RPCs not only confers competence to generate Müller glial and bipolar cells on late-stage RPCs, but also functions to promote cell cycle exit.

## DISCUSSION

This study represents the first large-scale cellular profile of gene expression changes across the full course of neurogenesis in any mammalian CNS region. The large number of individual cells profiled allows us to reconstruct a continuous representation of changes in gene expression that occur during the course of retinal neurogenesis and specification of the major retinal cell types. This represents a major advance beyond previous work and has broad implications for studies of neural development. Previous expression profiling studies of developing retina, even when supplemented with large-scale in situ hybridization, could not provide interpretable expression data at the cellular level (Blackshaw *et al.*, 2001; Blackshaw *et al.*, 2004; Hoshino *et al.*, 2017). Likewise, previous single cell gene expression profiling of retinal progenitors has been performed previously, but only on a few hundred cells using semi-quantitative microarray-based analysis (Trimarchi *et al.*, 2008). ScRNA-Seq studies of neurogenesis in brain regions such as cerebral cortex and midbrain have analyzed only a few thousand cells, usually at a relatively smaller number of developmental ages (La Manno *et al.*, 2016; Mayer *et al.*, 2018; Nowakowski *et al.*, 2017; Zhong *et al.*, 2018), and thus do not fully capture the transcriptional changes associated with the initiation of neurogenesis, across development competence, and loss of proliferative potential.

Previous studies have suggested that RPCs which express neurogenic bHLH factors such as Neurog2, Atoh7 and Olig2 may be intrinsically biased to generate specific neuronal subtypes, and show limited gliogenic competence (Brzezinski *et al.*, 2011; Brzezinski *et al.*, 2012; Hafler *et al.*, 2012). This study identifies a clear transcriptional distinction between the uncommitted population of RPCs, and a population of actively-proliferating cells expressing a set of genes that include neurogenic bHLH factors. Significant transcriptional differences are also observed between early and late stages in both bonafide and neurogenic RPCs, consistent with changes in RPC competence over development. As we observe most of the RPC transcriptome to be temporally dynamic, we suggest that the relative developmental age of an individual RPC can be determined using these data. Nonetheless, beyond the categories described here, we do not see evidence for molecularly distinct RPC subtypes at individual ages suggesting a more probabilistic model for cell fate specification within this system.

Since these data represent a snapshot in time for each individual cell, neither division mode or cell cycle duration can be directly assessed. However, the increase in cell cycle component variation within RPCs at individual ages as development progresses is suggestive of the increase in cell cycle length. Additionally, we observe a progressive increase in Notch signaling. These transitions may reflect changes that occur continuously over the course of neurogenesis, such as the shift between symmetric and asymmetric modes of cell division (Livesey and Cepko, 2001), and the increase in retinal cell cycle length (Alexiades and Cepko, 1996). In contrast, competence transitions appear to be more discrete with sharper distinctions between early versus late populations of bonafide and neurogenic RPCs cells. Further, transitions between neuroepithelial cells and proliferating RPCs are sharp, as are transitions between the main and neurogenic RPC population.

Both supervised pseudotemporal and unsupervised scCoGAPS analyses identified hundreds of candidates for genes that regulate developmental competence transitions and/or postmitotic differentiation of individual retinal cell types. Notably, many of these genes show highly dynamic and complex expression patterns, and they are expressed in discrete temporal windows in multiple cell types. Such complex expression patterns are more the rule than the exception, and they are readily identifiable from scCoGAPS patterns. As a consequence, functional analysis of such genes needs to be conducted and interpreted carefully in future studies. Whereas scCoGAPS finds patterns shared across cells, additional techniques are essential to quantify the relative heterogeneity of cells at specific developmental time points. Here, we apply the EVA statistic (Afsari *et al.*, 2014) to quantify differences in transcriptional heterogeneity of discrete gene sets as a measure of relative pathway dysregulation amongst cells that are aggregated by a common feature -- in our case, by developmental age.

The analysis described here allowed us to identify the NFI family of transcription factors as candidate regulators of temporal cell fate specification within the developing retina. Previous studies have demonstrated an essential role for *Nfia* in specification of astrocytes in several CNS regions (Deneen *et al.*, 2006; Glasgow *et al.*, 2014; Kang *et al.*, 2012), and we observe that loss of function of *Nfia, Nfib* or *Nfix* individually all lead to defects in Müller glial specification. However, analysis of *Nfia/b* double or *Nfia/ b/x* triple mutants indicate that these genes are necessary for generation of both Müller glia and late-born bipolar neurons. In addition to, and possibly a consequence of specifying glial and bipolar cell identity, these transcription factors are also necessary for cell cycle exit in late-stage RPCs. While *Nfix* has been previously reported to inhibit proliferation of both ES-derived neuroepithelial cells in culture (Martynoga *et al.*, 2013) and postmitotic neural stem cells of the SVZ (Heng *et al.*, 2015), *Nfia* was previously shown to promote astrocyte proliferation (Glasgow *et al.*, 2013). Surprisingly, the high levels of ongoing proliferation and retinal neurogenesis in mature animals observed here in *Nfia/b/x* mutants has not been previously reported. This dual role in regulating both proliferative quiescence and developmental competence has not been previously reported in the developing CNS, but may represent a widely used mechanism for generating appropriate numbers of late-born cells. Finally, since *Nfia, Nfib* and *Nfix* are all expressed in mature Müller glia (de Melo *et al.*, 2016a) (Figure 5), this raises the tantalizing possibility that targeted inactivation of these genes may contribute to induction of both proliferation and neurogenic competence in these cells, and so prove useful in regenerative therapies that seek to replace dead or dying photoreceptors in blinding disorders.

With few exceptions, the cell diversity of other CNS regions are considerably greater, and their development far less well characterized. Unbiased high-throughput analysis of gene expression at the single cell level, in conjunction with both unsupervised pattern identification and supervised pseudotime analysis like that described here, will ultimately provide the potential to understand the gene regulatory networks that give rise to the immense diversity of cell types within the rest of the CNS.

## ACKNOWLEDGEMENTS

This work was supported by grants from the NIH (R01EY020560 and U01EY027267 to SB, F32EY024201 and K99EY027844 to BSC, K08EY027093 to FR, R01CA177669 and U01CA212007 to EJF), NYSTEM awards C026429, C030133 and C30290GG (RMG), the Chan-Zuckerberg Initiative DAF (2018-183445 to LAG, and 2018-183444 to EJF), the Johns Hopkins University Catalyst (EF & LAG) and Discovery awards (EJF), and the Johns Hopkins University School of Medicine Synergy Award (SB, LAG, & EJF). The authors would like to thank C.A. Berlinicke and D.J. Zack for assistance with FACS analysis, the Johns Hopkins P30 Center for Neuroscience Research (NS050274), use of the 10x Genomics Single Cell system through the Genetic Resources Core Facility; the Johns Hopkins Institute of Genetic Medicine, Baltimore, MD, and the Hopkins Microarray and Deep Sequencing Core for assistance with sequencing; and J. Nathans, A. Kolodkin, R. Bremner, W. Yap, A. McCallion, and D.W. Kim for comments on the manuscript.

## AUTHOR CONTRIBUTIONS

BSC, LAG, and SB conceived the study. BSC, GSO’B, EJF, LAG and SB directed the study. BSC generated scRNA-Seq data, with assistance from GHC. GSO’B, BSC, TS, ED, LAG and EJF analyzed scRNA-Seq data, with LAG & EJF as senior bioinformaticians. GSO’B, TS, LAG, and EJF developed scCoGAPS. ED, LAG, and EJF extended EVA for scRNA-Seq data. BSC and FS analyzed the function of NFI factors in mouse retina, with assistance from FR and REJ-E. RMG provided Nfia/ b/x-floxed mice. BSC, GSO’B, EJF, LAG, and SB wrote the paper, with input from all co-authors.

## DATA AVAILABILITY

Single cell RNA-Seq count data are available for direct download at https://github.com/gofflab/developing_mouse_retina_scRNASeq. Raw sequencing data is deposited with the NCBI Short Read Archive and Gene Expression Omnibus under acession number XXXXXXX.

## METHODS

### Animals

Timed pregnant CD-1 mice used for droplet-based single cell RNA sequencing, in situ hybridization, and electroporation were purchased from Charles River Laboratories. For Smart-Seq2 analysis, Chx10-Cre-GFP mice were used (Rowan and Cepko, 2004). *Nfia^lox/lox^; Nfia^lox/lox^*, and *Nfix^lox/lox^* mice were crossed to *Chx10*-Cre-GFP to generate RPC-specific loss of function mutants of these genes. Mice were housed in a climate-controlled pathogen free facility, on a 14 hour-10 hour light/dark cycle (07:00 lights on-21:00 lights off). All experimental procedures were preapproved by the Institutional Animal Care and Use Committee of the Johns Hopkins University School of Medicine.

### Tissue Dissociation

Eyes were enucleated from animals and retinas dissected in fresh and cold 1X PBS, using eyes from approximately one litter of animals for each sample in order to ensure appropriate numbers of cells were captured for downstream analyses. Dissected retinas were then transferred to 200μl of cold HBSS per retina (P14) or an approximate equivalent amount of tissue for younger ages. An equivalent amount of Papain solution (for 1ml - 700μl reagent grade water, 100μl of freshly prepared 50mM L-Cysteine (Sigma), 100μl 10mM EDTA, 10μl 60mM 2-mercaptoethanol (Sigma), and Papain added to 1mg/ml (Worthington)) was added and incubated at 37°C for 10 minutes, with slight trituration performed every 2 minutes. 600μl of Neurobasal Media supplemented with 10% FBS was added for every 400μl of dissociation solution, and samples were further dissociated with gentle pipetting. Samples were subjected to DNAse treatment (5μl DNAseI (RNAse free Recombinant DNAseI; Roche) for every 1ml of dissociation solution; 5 minutes at 37°C). Cells were then pelleted through centrifugation for 5 minutes at 300 RCF. Liquid was carefully aspirated off the cell pellet, followed by resuspension of the pellet in 1-5ml Neurobasal media with 1% FBS, depending on required concentration of cells in suspension. Cellular aggregates were removed by straining cells through a 50μm filter.

### Single cell library preparation

Smart-Seq2 analysis was performed on individual sorted Chx10-Cre-GFP (+) RPCs isolated by FACS into 96-well plates, and processed as previously described (Chevee *et al.*,2018). Single cell suspensions for 10x libraries were loaded onto the 10x Genomics Chromium Single Cell system using the v2 chemistry per manufacturer’s instructions (Zheng *et al.*, 2017). Approximately 17,000 live cells were loaded per sample in order to capture transcripts from roughly 10,000 cells. Estimations of cellular concentration and live cells in suspension was made through Trypan Blue staining and use of the Countess II cell counter (ThermoFisher). Single cell RNA capture and library preparations were performed according to manufacturer’s instructions. Sample libraries were sequenced on the NextSeq 500 (Illumina).

### Library preprocessing

Sequencing output was processed through the Cell Ranger 1.2.1 mkfastq and count pipelines using default parameters. Reads were quantified using the mouse reference index provided by 10x genomics (refdata-cellranger-mm10 v1.2.0). Raw count matrices for individual runs were manually aggregated and cells were given unique, sample-specific cell identifiers to prevent duplication of nonunique barcodes across samples. The full raw count matrix was then used as input for the Monocle2 single cell RNA-Seq framework (Trapnell *et al.*, 2014).

### Coarse assignment of cell type at individual time points

tSNE-dimension reduction was performed on the top principal components learned from high variance genes in cells captured at individual timepoints. Mclust version 5.4 (Scrucca *et al.*, 2016) was used to cluster cells in tSNE-space at which point cell type identity of clusters was assigned based on expression of known marker genes for either retinal or non-retinal tissue.

### Cell normalization, identification of high variance genes, and differential testing

After coarse annotation of cells from individual time points, the 10x data were manually aggregated to create a comprehensive dataset. Initial cell type designation was used to aid in supervising downstream analyses. To identify genes with higher variation than expected, we first normalized for sequencing depth using the Waddington-OT transformation to transcript copies per 10,000 (CPT) (Schiebinger *et al.*, 2017). To identify high-variance genes, a generalized additive model (MGCV R package (Wood, 2011)) was fit to the log2 mean CPT versus a cubic spline fit to the log2 coefficient of variation (BCV) across all genes with detectable expression in at least 5 cells (Figure S3). Genes with residuals to this fit greater than or equal to 1.5 were chosen as ‘high-variance’ genes and the log2 CPT or CPC values for these selected genes were used as input for downstream analyses as appropriate. All differential expression tests were performed across all expressed genes using the Monocle2 VGAM likelihood ratio test (Trapnell *et al.*, 2014). In all cases, the number of genes detected in each cell was included in both the full and reduced models as a nuisance parameter.

### Visualization of global cellular state(s)

Dimensionality reduction and visualization for the aggregate 10x data was performed using Uniform Manifold Approximation and Projection (UMAP) (McInnes and Healy, 2018) for all cells passing QC. Briefly the first 20 principle components of the log10(CPT+1) of the high-variance genes was used as input for the python implementation of the UMAP algorithm with the following additional parameters: min_dist = 0.3, n_neighbors=30, random_state=1, n_components=3, metric=“canberra”. The resulting 3 dimensional embedding was imported into R and visualized using the rgl package.

### Clustering analysis and final cell type assignment

Clustering on the UMAP embedding was performed using k-means clustering on UMAP coordinates. Cell type assignment was informed by both marker gene expression and previous coarse cell type annotation from clustering performed on the individual ages. Clustering analysis of the amacrine trajectory to delineate amacrine and horizontal cells was performed using k-means clustering on the largeVis coordinates.

### Pseudotemporal analysis

Monocle pseudotemporal analysis was performed on the high variance gene set derived from the subsets of cells being analyzed, altering the dimension parameter to refine resulting trees to reflect both known biology and terminal states comprised largely of single cell types. For pseudotemporal analysis performed on the subset of RPCs, the following additional parameters were used during dimension reduction: tol =1.0e-8, lambda=400*ncol(CellDataSet). The root state was identified as the state that contained the majority of cells with the earliest developmental age for each individual analysis. Genes with significant expression changes as a function of pseudotime were identified using the Monocle differential gene test, using a multiple-testing corrected q-value cutoff of 1.0e-5. Cell type identity of individual pseudotime states were assigned based on the cell type identity of the majority of cells within a given branch. BEAM tests were performed on most major branch points of the cellular hierarchy using all default parameters with the exception of the dimensionality of the embedding.

### Pattern discovery

scCoGAPS and PatternMarker analysis was performed using the R/Bioconductor package CoGAPS version 3.0.0 as described (Stein-O’Brien *et al.*, in prep). Briefly, the expression matrix of high variance genes was subset into 200 sets of cells for parallelization. A sampling scheme, using expertly curated cell type annotations, was used in order to ensure representation of rare cell types in each set. A static ratio of cell types was established to reflect biological prevalence and diversity of each cell type while allowing for adequate representation. Cells were then sampled with replacement to ensure the necessary numbers to maintain this ratio in all sets. The resulting 200 sets of 1500 cells each were then run in parallel over a range of dimensionalizations. Consensus amplitude signatures were derived by a matching algorithm designed to ensure robustness of signatures (Star Methods). Pattern weights for all the cells were then learned in parallel from these signatures to ensure reciprocity across all of the sets. The PatternMarker statistic was calculated as previously described (Stein-O’Brien *et al.*, 2017b). The entire pipeline has been compiled into the scCoGAPS function found within the CoGAPS package starting at version 3.0.0.

### Pathway dysregulation analysis

We quantify pathway dysregulation using EVA from the R/Bioconductor package GSReg version 1.17.0 (Afsari *et al.*, 2014). Briefly, EVA computes the Kendal-tau dissimilarity between transcriptional profiles of genes in a pathway for all cells in one group and compares their expected dissimilarity to that computed for all the cells in another group using U-theory statistics. This statistic quantifies relative pathway dysregulation between cells in these two conditions. Because the Kendal-tau dissimilarity is rank-based, it is robust to normalization and read depth, but ill-defined for missing values. To address this, we imputed scRNA-Seq data from RPCs with MAGIC version 0.1.0 (Python) prior to analysis. EVA is applied to gene sets for the cell cycle from GeneGlobe pathways, FGF pathway from the reactome pathway database, NOTCH pathway from the hallmark gene sets, and WNT pathway from the hallmark gene sets.

### in situ hybridization

Whole heads (E12-P5) or enucleated eyes of animals were placed directly into Tissue-Tek OCT media (VWR) and frozen and stored at −80°C prior to sectioning. Section RNA in situ hybridization was performed as previously described (Blackshaw *et al.*, 2004).

### Immunohistochemistry and EdU staining

Eyes were enucleated from animals and placed in cold 4% paraformaldehyde for 1 hour. Retinas were then dissected and placed into 30% sucrose in PBS overnight at 4°C, after which they were mounted in Tissue-Tek OCT media (VWR) and frozen prior to sectioning. Immunohistochemistry was performed as previously described (de Melo *et al.*, 2016a). Antibodies utilized for fluorescent immunohistochemistry are as follows: goat anti-Brn3 (1:200; Santa Cruz Biotechnology; SC-6026 (C-13)), mouse anti-calbindin-D-28K (Calb1) (1:200; Sigma-Aldrich; C9848 (clone CB-955)), sheep anti-Chx10 (Vsx2) (1:500; Exalpha Biologicals; X1180P), rabbit anti-GFP (1:1000; Invitrogen; A6455), mouse anti-Glutamine synthetase (Glul) (1:200; BD Biosciences; 610518 (Clone6)), mouse anti-Islet1 (1:200; Developmental Studies Hybridoma Bank; 40.2D6), mouse anti-Ki67 (1:200; BD Biosciences; 550609 (Clone B56)), rabbit anti-Lhx2 (1:1000; generated in house with Covance), mouse anti-P27Kip1 (1:500; BD Transduction Labs; 610241 (clone57/Kip1/p27)), mouse anti-Pax6 (1:200; Developmental Studies Hybridoma Bank; Pax6a.a 1-223), mouse anti-Nfia/b (1:200; CDI Labs; R1356.1.2C6 (Venkataraman *et al.*, 2018)), mouse anti-Tfap2a (1:200; Abnova; Clone 2G5), rabbit anti-Recoverin (1:200; Millipore; Lot LV1480447), mouse anti-Pkcα (1:200; Millipore; Clone M4), rabbit anti-GFAP (1:500; DakoCytomation; Z0334), mouse anti-Rho4D2 (Rhodopsin) (1:200, Gift from Bob Molday), rabbit anti-- -catenin (Ctnnb1) (1:200; Sigma; C2206), and rabbit anti-Iba1 (1:400, Wako; 019-19741).

Secondary antibodies used were donkey anti-rabbit 488 (1:500; Jackson ImmunoResearch), goat anti-rabbit 555 (1:500; Invitrogen), donkey anti-Rabbit 594 (1:500, Invitrogen), goat anti-rabbit 633 (1:500; Invitrogen), donkey anti-rabbit 647 (1:500; Jackson ImmunoResearch), donkey anti-mouse 555 (1:500; Invitrogen), goat antimouse 555 (1:500; Invitrogen), donkey anti-mouse 594 (1:500; Jackson ImmunoResearch), donkey anti-mouse 647 (1:500; Jackson ImmunoResearch), donkey antigoat 594 (1:500; Invitrogen), donkey anti-goat 633 (1:500; Invitrogen), donkey anti-sheep 568 (1:500; Invitrogen), donkey anti-sheep 647 (1:500; Jackson ImmunoResearch).

For EdU analyses, animals were injected with 50 mg/kg EdU (10 mM in saline) at P21 and euthanized at P28. EdU staining was performed using the Click-IT EdU AlexaFluor 647 imaging kit (Invitrogen), with slides placed into blocking steps for the immunohistochemistry protocol directly after EdU detection. Nuclei were counterstained with DAPI (1:5000) and coverslipped using Vectashield (Vector Labs). Immunohistochemical data shown was imaged and photographed on either the BZ-X700 microscope (Keyence) or using a LSM 700 confocal (Zeiss).

## SUPPLEMENTAL FIGURE LEGENDS

**Figure S1. Smart-Seq2 dataset.** (A) Density plot of total mRNAs per cell after Monocle relative2abs transformation. (B) Tables of number of cells per plate (top), number of cells at each developmental age (middle), and QC metrics of the entire Smart-Seq2 dataset. (C) Graph of the number of genes expressed per cell, faceted by developmental age. (D) Dispersion plot of the high variance genes (red) used for dimension reduction. Black line indicates gam fit. E) tSNE dimension reduction with cells colored by the plate that they came from. F-I) Cellular heatmaps of reduced tSNE maps for (F) markers of proliferative RPCs (Ccnd1, Cdk4, Dkk3, Pax6), (G) markers of the G1/S phases of the cell cycle (Ccne2 and Pcna) or G2/M (Ccnb1 and Ube2c), (H) markers of neurogenic progenitors (Atoh7, Neurod1, Neurog2, and Olig2), or (I) cell type-specific markers including those of photoreceptors (Crx), retinal ganglion cells (Isl1 and Pou4f2) and amacrine cells (Tfap2b).

**Figure S2. 10x Dataset.** (A) Number of replicate datasets and total number of single cells profiled corresponding to each developmental age. (B) Breakdown of QC metrics according to the individual samples, including number of cells, mean reads per cell, and total number of sequencing reads of each library. (C) QC metrics of the unique molecular identifiers (UMI) and Genes detected across all the cells. D-E) Boxplots of the total mRNAs or UMIs detected per cell faceted by (D) developmental age or (E) sample. F-I) UMAP dimension reduction of the 10x dataset with cells colored by (F) developmental age, (G) the original cell type annotation based on clustering from individual ages, (H) cluster designation following Mclust analysis, (I) or re-annotation of Mclust clusters in UMAP space. Re-annotation of cell type was based on expression of known marker genes of retinal cell types within each cluster. (J) Graph displaying the proportion of retinal cell type profiled at each developmental age. (K) Graph displaying the proportion of each developmental age that each retinal cell type was captured. (L) Graph displaying the proportion of mature retinal cell types captured at each developmental age with RPCs and neurogenic cells eliminated from the analysis. (M) Graph displaying the proportion of RPC, neurogenic and gliogenic (or mature Müller glia) cells capture at each developmental age.

**Figure S3. Dispersion plots.** (A) Dispersion plots displaying high variance genes (black) across all cells profiled from individual ages. (B-C) Dispersion plots of high variance genes across aggregated cells across the (B) entire dataset and used for UMAP dimension reduction, or (C) annotated retinal cells and used for developmental order in pseudotime analysis across the retinal cells.

**Figure S4. tSNE plots of 10x Samples by developmental age.** tSNE dimension reduction of 10x datasets across all cells at (A) E11, (C) E12, (E) E14, (G) E16, (I) E18, (K) P0, (M) P2, (O) P5, (Q) P8, (S) P14 and annotated retinal cells at (B) E11, (D) E12, (F) E14, (H) E16, (J) E18, (L) P0, (N) P2, (P) P5, (R) P8, (T) P14. Cells are colored by annotated cell type after clustering performed on tSNE plots of individual samples. Cell types were re-annotated after samples were subset to retinal cells only and recursive tSNE dimension reduction was performed on a new set of high variance genes within the retinal cells.

**Figure S5. Amacrine and horizontal cell trajectory.** (A) LargeVis dimension reduction on the subset of cells from UMAP representations annotate as amacrine cells. Colors represent re-annotation of cell types based on known marker gene expression within clusters of cells. (B) UMAP representation indicating the position of the starburst amacrine cells and horizontal cells within the full retina dataset. Starburst amacrine cells are classified as amacrine cells within Figure 1G and subsequent analyses on the full dataset. (C) Complex pseudotime tree on the amacrine cell trajectory with cells colored by annotated cell type from the largeVis dimension reduction. (D) Examples of gene expression across pseudotime of transcripts that display differential expression across the annotate amacrine, horizontal and starburst amacrine cells. Transcript expression within aggregated cell types across pseudotime is represented by the alternately colored lines. Abbreviations: Photo. Precurs. - Photoreceptor Precursors; RGCs - Retinal Ganglion Cells.

**Figure S6. Complex pseudotime trees of annotated cell features.** (A-C) Complex pseudotime trees resulting from pseudotime analysis on ~32,000 retinal cells and colored by (A) Pseudotime value, (B) developmental age, and (C) annotated cell type. C) Complex pseudotime trees are faceted to show cells with individual cell type annotations.

**Figure S7. BEAM analysis - bipolar and photoreceptor cells versus amacrine/horizontal cells.** Heatmap of transcripts that display state-dependent differential expression between the bipolar and photoreceptor cell branch (red in complex pseudotime tree) and the amacrine/horizontal cell branch. Heatmaps represent gene expression across pseudotime starting from the top of the complex pseudotime tree (center of heatmaps) and progressing through the pseudotime tree towards the celltypes of interest (periphery of heatmaps). All genes displayed significant differential expression (q < 1.0e-20) by the BEAM test.

**Figure S8. BEAM analysis - bipolar cells versus photoreceptor cells.** Heatmap of transcripts that display state-dependent differential expression between the bipolar (red in complex pseudotime tree) and the photoreceptor cell branch (blue in complex pseudotime tree). Heatmaps represent gene expression across pseudotime starting from the top of the complex pseudotime tree (center of heatmaps) and progressing through the pseudotime tree towards the cell types of interest (periphery of heat-maps). All genes displayed significant differential expression (q < 1.0e-20) by the BEAM test.

**Figure S9. BEAM analysis - late cells versus early cells.** Heatmap of transcripts that display state-dependent differential expression between the late cell types (red in complex pseudotime tree) and the early cell types (RGCs and early RPCs; blue in complex pseudotime tree). Heatmaps represent gene expression across pseudotime starting from the top of the complex pseudotime tree (center of heatmaps) and progressing through the pseudotime tree towards the cell types of interest (periphery of heat-maps). All genes displayed significant differential expression (q < 1.0e-20) by the BEAM test.

**Figure S10. BEAM analysis - RGCs versus early RPCs.** Heatmap of transcripts that display state-dependent differential expression between the RGCs (red in complex pseudotime tree) and the Early RPCs (blue in complex pseudotime tree). Heatmaps represent gene expression across pseudotime starting from the top of the complex pseudotime tree (center of heatmaps) and progressing through the pseudotime tree towards the celltypes of interest (periphery of heatmaps). All genes displayed significant differential expression (q < 1.0e-20) by the BEAM test.

**Figure S11. BEAM analysis - Late RPCs and Müller glia versus amacrine/horizontal, bipolar and photoreceptor cells.** Heatmap of transcripts that display state-dependent differential expression between the late RPCs and Müller glia (red in complex pseudotime tree) and the amacrine/horizontal cells, bipolar cells and photoreceptors (blue in complex pseudotime tree). Heatmaps represent gene expression across pseudotime starting from the top of the complex pseudotime tree (center of heatmaps) and progressing through the pseudotime tree towards the cell types of interest (periphery of heatmaps). All genes displayed significant differential expression (q < 1.0e-20) by the BEAM test.

**Figure S12. BEAM analysis - Late RPCs versus Müller glia.** Heatmap of transcripts that display state-dependent differential expression between the late RPCs (red in complex pseudotime tree) and the Müller Glia (blue in complex pseudotime tree). Heatmaps represent gene expression across pseudotime starting from the top of the complex pseudotime tree (center of heatmaps) and progressing through the pseudotime tree towards the cell types of interest (periphery of heatmaps). All genes displayed significant differential expression (q < 1.0e-20) by the BEAM test.

**Figure S13. BEAM analysis on amacrine Cell trajectory pseudotime analysis - starburst Amacrine versus other amacrine cells.** Heat-map of transcripts that display state-dependent differential expression between the Starburst amacrine cells (red in complex pseudotime tree) and other amacrine cells (blue in complex pseudotime tree). Heatmaps represent gene expression across pseudotime starting from the top of the complex pseudotime tree (center of heatmaps) and progressing through the pseudotime tree towards the celltypes of interest (periphery of heatmaps). All genes displayed significant differential expression (q < 1.0e-20) by the BEAM test.

**Figure S14. BEAM analysis on amacrine cell trajectory pseudotime analysis - horizontal Cells versus amacrine cells.** Heatmap of transcripts that display state-dependent differential expression between the horizontal cells (red in complex pseudotime tree) and amacrine cells (blue in complex pseudotime tree). Heatmaps represent gene expression across pseudotime starting from the top of the complex pseudotime tree (center of heatmaps) and progressing through the pseudotime tree towards the celltypes of interest (periphery of heatmaps). All genes displayed significant differential expression (q < 1.0e-20) by the BEAM test.

**Figure S15. Bonafide RPC and neurogenic RPC cell pseudotime.** (A) Heatmap of the normalized expression of 40 core Notch-pathway genes, as identified from MSigDB, across pseudotime. Cells were grouped into 11 groups encompassing the entire range of the RPC pseudotemporal analysis from early pseudotime (top rows of heatmap) through late pseudotime (bottom rows of heatmap) (B) Heatmap of differentially expressed genes across pseudotime of the subset of RPCs and grouped by developmental age. (C) Results of EVA analysis displaying relative dissimilarity of the Fgf- (top), Notch- (middle), and Wnt-signaling pathways within RPCs from E11 to P5. D) Graph of the proportions of cells arising from various developmental ages across the RPC, Neurogenic Cell, and Müller glial cell pseudotime analysis. (E) Pseudotime density plot of cells in the RPC, Neurogenic Cell, and Müller glial cell pseudotime analysis colored by developmental age, showing three relative clusters corresponding to neuroepithelial cells, early RPCs and neurogenic cells, and late RPCs, late neurogenic cells and Müller glial cells. (F) Graph depicting the proportions of cells at individual developing ages in the RPC, Neurogenic Cell, and Müller glial cell pseudotime analysis that are annotated as individual cell types. *** indicates p<0.001 from EVA analysis; ns - not significant

**Figure S16. scCoGAPS pattern weights within retinal cells.** UMAP dimension reduction plots of all retinal cells, colored by scCoGAPS pattern weights. Individual cell pattern weights are plotted for each cell, with corresponding values progressing from low pattern weight (grey) to high pattern weight (blue).

**Figure S17. scCoGAPS pattern weights within complex pseudotime plots of retinal cells.** Complex pseudotime trees displaying pattern weight of individual cells within pseudotime on the subset of ~32,000 retinal cells. Individual cell pattern weights are plotted for each cell, with corresponding values progressing from low pattern weight (grey) to high pattern weight (blue).

**Figure S18. NFI transcription factors expression within UMAP dimension reductions.** (A-C) UMAP dimension reduction plots displaying expression of (A) Nfia, (B) Nfib, and (C) Nfix within RPCs, displaying enriched expression of the NFI transcription factors within late RPCs. D-F) UMAP dimension reduction plots displaying expression of (D) Nfia, (E) Nfib, and (F) Nfix within the entire retina dataset. Expression of each NFI transcription factor is observed in late RPCs, a fraction of cells within the amacrine cell trajectory and within the Müller glial cell trajectory. Cellular expression is colored on a scale of low (grey) to high (red) expression.

**Figure S19. Immunohistochemistry of cell type markers within NFI transcription factor conditional knockout retinas.** Immunohistochemistry performed on P14 retinas examining changes in marker gene protein expression across control or Chx10:Cre-GFP mediated conditional knockouts of Nfia, Nfib, Nfix, Nfia/b, or Nfia/b/x. Chx10:Cre-GFP transgene is shown in green, with all nuclei counterstained in merged images. (A) Immunohistochemistry for NFI protein expression shows efficiency of NFI transcription factor knockout. B-D) Retinal progenitor cell markers (B) Lhx2, (C) Pax6 and (D) Vsx2 are not up-regulated with the maintenance of RPC proliferation in Nfia/b and Nfia/b/x conditional knockouts. (B) Lhx2-positive Müller glia are lost in Nfia/b and Nfia/b/x conditional knockouts. (C) Amacrine cell marker Pax6 is maintained D) A significant fraction of Vsx2-positive bipolar cells are lost in Nfia/b/x triple mutants. (E) Expression of Brn3-positive retinal ganglion cells. (F) Expression of Calbindin-positive horizontal cells. (G) Expression of Tfap2a-positive amacrine cells. (H) Most of the Isl1-positive bipolar cells are lost in Nfia/b/x triple conditional mutant retinas. (I) Expression of Recoverin-positive photoreceptors. (J) Both Nfia/b and Nfia/b/x conditional mutants show a large loss of P27Kip-positive Müller glial cells. Asterisk indicate regions in which cell type markers all lost. Arrowheads indicate remaining sparse labeling of cell type markers. Scale bars - 100μm

**Figure S20. Conditional knockout of NFI transcription factors results in loss of retinal architecture, glial activation and maintenance of RPC proliferation.** A) Immunohistochemistry on P14 retinas for glial fibrillary activating protein (GFAP), marking the vasculature overlaying the nerve fiber layer (asterisk) and activated Müller glia (arrowheads in Nfia/b and Nfia/b/x conditional mutants). (B) Expression of beta-catenin (Ctnnb1) in P14 retinas marks the localization of tight junctions. Loss and disruption of b-catenin expression within the OLM (arrowheads) is indicative of Müller glial cell loss. (C) Resident or activate microglia are marker by Iba1 expression in P14 retinas. (D) Quantification of the distance from the outermost edge of inner plexiform layer (IPL) to the outer limiting membrane (OLM) or edge of the nuclear layer when no OLM remains. p < 0.0001. (E) Quantification of the number of microglia present per P14 retinal section in control and NFI transcription factor conditional mutants retinas. p < 0.0001. (F) Quantification of the number of proliferative cells in P28 retinas. EdU pulse was given at P21 and chased until P28. P28 retinas were stained with proliferative cell marker Mki67. (G) Quantification of the percentage of EdU positive cells that co-labeled with either Mki67 or Recoverin in Nfia/b/x triple mutant retinas. Statistics are the results of a one-way ANOVA (D and E). Tukey’s multiple comparisons tests results are indicated as * - p < 0.05 ** - p < 0.01, *** - p < 0.001, and **** - p < 0.0001 in panels D and E. Statistics in panel F represent results of unpaired t-test, **** - p < 0.0001. n >= 3 animals per condition. For distance measurements in (D), 3 measurements per retinal section were averaged per animal to account for potential section plane and localization (central to peripheral retina) differences that could affect distance metrics. Scale bars - 100μm

### Movies

Movie 1 – UMAP-dimension reduction of droplet-based single cell RNA-sequencing of single developing mouse retinal cells with samples colored by developmental age. Related to Figure S2F.

Movie 2 – UMAP-dimension reduction of droplet-based single cell RNA-sequencing of single developing mouse retinal cells with samples colored by annotated cell type as determined by marker gene expression in clustered cells. Related to Figure S2I.

Movie 3 – UMAP-dimension reduction of droplet-based single cell RNA-sequencing of single developing mouse retinal cells with samples colored by annotated cell type as determined by marker gene expression in clustered cells. Extra-retinal and doublet cells have been removed. Related to Figure 1F.

Movie 4 – UMAP-dimension reduction of retinal cells colored by sc-CoGAPS Pattern 36 (sex pattern) pattern weights. Cells are colored on a scale from gray-blue corresponding to low-high pattern weight. Related to Figure 4C.

Movie 5 – UMAP-dimension reduction of retinal cells colored by sc-CoGAPS patterns corresponding to cell cycle phase. Cells are colored by pattern weights from low (gray) to high (pattern color) with color mixing highlighting contributions of multiple patterns within the same cell. Related to Figure 4D.

Movie 6 – UMAP-dimension reduction of retinal cells colored by combinations of pattern weights for scCoGAPS patterns corresponding to neurogenic commitment (red) and photoreceptor specification (green) through rod differentiation (blue). Cells are colored by pattern weights from low (gray) to high (pattern color) with color mixing highlighting contributions of multiple patterns within the same cell. Related to Figure 4E.

## SUPPLEMENTAL TABLES

Table S1 - Smart-Seq2 high variance genes. Related to Figure 1B-D.

Table S2 - Smart-Seq2 differential gene test - RPCs. Related to Figure 1B-D.

Table S3 - Smart-Seq2 differential gene test - All cell types. Related to Figure 1B-D.

Table S4 - High variance genes used for UMAP dimension reduction on 10x samples. Related to Figure 1E-F and Figure S2F-I.

Table S5 - 10x cell type features. Related to Figure 1E-G and Figure S2J-M.

Table S6 - High variance genes across 10x retinal cells. Related to Figure 2.

Table S7 - Genes displaying differential expression across retina pseudotime. Related to Figure 2D.

Table S8 - Pattern markers from scCoGAPS on 10x samples. Related to Figure 4.

Table S9 - Gene weights within scCoGAPS patterns. Related to Figure 4.

